# Sex-specific effects of fecal microbiota transplantation on TBI-exacerbated Alzheimer’s pathology in mice

**DOI:** 10.1101/2025.09.11.675717

**Authors:** Sirena Soriano, Austin Marshall, Morgan Holcomb, Hannah Flinn, Marissa Burke, Göknur Kara, Paula Scalzo, Sonia Villapol

## Abstract

**Background:** Traumatic brain injury (TBI) accelerates Alzheimer’s disease (AD) pathology and neuroinflammation, potentially via gut-brain axis disruptions. Whether restoring gut microbial homeostasis mitigates TBI-exacerbated AD features remains unclear, particularly with respect to sex differences.

**Objective:** The goal of our study was to test whether fecal microbiota transplantation (FMT) modifies amyloid pathology, neuroinflammation, gut microbial composition, metabolites, and motor outcomes in male and female 5xFAD mice subjected to TBI.

**Methods:** Male and female 5xFAD mice received sham treatments or controlled cortical impact, followed 24 hours later by vehicle (VH) or sex-matched FMT from C57BL/6 donors. Assessments at baseline, 1, and 3 days post-injury included Thioflavin-S and 6E10 immunostaining for Aβ, Iba-1 and GFAP for glial activation, lesion volume, rotarod performance, 16S rRNA sequencing for microbiome profiling, serum short-chain fatty acids (SCFAs), and gut histology.

**Results:** TBI increased cortical and dentate gyrus Aβ burden, with females showing greater vulnerability. FMT reduced Aβ deposition in sham animals and shifted plaque morphology but did not attenuate TBI-induced amyloid escalation. FMT differentially modulated glial responses by sex and region (reduced microgliosis in males) without altering lesion volume. Rotarod performance was better in sham females compared to males and declined in FMT-treated TBI females. Fecal microbiome alpha diversity and richness were unchanged, while beta diversity revealed marked, time-dependent community shifts after TBI that were slightly altered by FMT. Gut morphology remained broadly intact, but crypt width increased after TBI, particularly in males.

**Conclusion:** In 5xFAD mice, TBI drives sex-dependent worsening of amyloid pathology, neuroinflammation, and dysbiosis. Acute FMT partially restores microbial composition and plaque features in sham animals but fails to reverse TBI-induced neuroinflammation or motor deficits. These findings underscore the context- and sex-dependence of microbiome interventions and support longer-term, sex-specific strategies for AD with comorbid TBI.

## 1. Introduction

Alzheimer’s disease (AD) is a progressive and debilitating neurodegenerative condition characterized by memory loss, cognitive decline, and pathological accumulation of amyloid-beta (Aβ) plaques and neurofibrillary tangles in the brain (DeTure and Dickson, 2019). Affecting millions of individuals globally, AD currently lacks effective disease-modifying therapies. A growing body of evidence suggests that the pathogenesis of AD is multifactorial, involving a complex interplay between genetic predisposition, environmental influences, neuroinflammation, and systemic metabolic dysfunction. Among emerging risk factors, traumatic brain injury (TBI) has gained considerable attention for its role in accelerating AD pathology and precipitating cognitive impairment (Livingston et al., 2024). TBI induces a cascade of neuropathological events, including neuronal injury, oxidative stress, blood-brain barrier (BBB) breakdown, and a sustained neuroinflammatory response (Sivandzade et al., 2020). These events are thought to facilitate Aβ deposition and tau hyperphosphorylation, creating a pro-AD environment in vulnerable individuals.

Recent studies have linked TBI to disruptions in the gut microbiota, an ecosystem of trillions of microorganisms that interact with the central nervous system through the gut-brain axis (El Baassiri et al., 2024). In our previous study, we found that within 24 hours of TBI, there was a marked shift in microbial composition, particularly among *Lactobacillus* strains (Treangen et al., 2018). Such dysbiosis has been reported in both TBI (Nicholson et al., 2019; Celorrio et al., 2021) and AD models (Harach et al., 2017; Vogt et al., 2017; Heston et al., 2023), where it is thought to exacerbate systemic inflammation and cognitive impairment (Zhan et al., 2018). The gut microbiota regulates the immune system, produces neuroactive metabolites such as short-chain fatty acids (SCFAs), and preserves epithelial barrier integrity (Parada Venegas et al., 2019). In our previous studies, administration of SCFA-producing probiotics after TBI reduced neuropathology (Holcomb et al., 2025), while acute-phase antibiotic treatment eliminated detrimental bacteria and conferred neuroprotection (Flinn et al., 2024). In AD, microbiota disruption has also been linked to increased intestinal permeability (“leaky gut”), elevated circulating proinflammatory cytokines, and heightened microglial activation, all of which contribute to neurodegeneration and cognitive decline (Popescu et al., 2024). These findings highlight modulation of the gut microbiota as a promising therapeutic strategy for neurodegenerative diseases.

Fecal microbiota transplantation (FMT) involves transferring intestinal microbiota from a healthy donor to a recipient with dysbiosis. Originally developed to treat recurrent *Clostridioides difficile* infection, FMT has since shown therapeutic potential in metabolic syndrome, inflammatory bowel disease, and neurodegenerative disorders (Kelly et al., 2016; Vendrik et al., 2020). In preclinical AD models, FMT has been reported to reduce Aβ pathology, improve memory function, and modulate neuroinflammation (Sun et al., 2019). In addition, our previous work demonstrated that young wild-type mice subjected to TBI and subsequently receiving FMT from AD animals exhibited exacerbated neuroinflammation and increased lesion size, indicating that the dysbiotic microbiome of AD donors can worsen post-TBI recovery (Soriano et al., 2022).

Despite these findings, the effects of FMT on TBI-exacerbated AD pathology, particularly concerning sex-specific responses, remain unclear. To address this gap, we employed the 5xFAD mouse model of AD, which develops early and aggressive Aβ plaque deposition at 5 months old. We subjected these mice to TBI to accelerate neuropathological progression. We hypothesized that FMT from young (2 months old), healthy wild-type donors could ameliorate TBI-induced gut dysbiosis and neuropathology, thereby restoring microbial homeostasis. In addition, we examined sex as a biological variable influencing treatment response. Our results support that TBI exacerbates neuroinflammatory and neuropathological outcomes in AD mice, with females exhibiting more pronounced pathology. Importantly, although FMT transiently restored aspects of microbial balance, it only reversed the accelerated Aβ accumulation of 5xFAD mice without injury. These findings underscore the role of FMT in driving sex-dependent neurodegenerative processes and highlight the need for more targeted microbiome-based interventions.

## 2. Material and methods

### 2.1 TBI in an AD mouse model

Young adult (2-month-old) C57BL/6J mice (Jackson Laboratories, Bar Harbor, ME) and 5xFAD (5-month-old) mice were housed at the Houston Methodist Research Institute animal facilities under a standard 12-hour light and dark cycle with ad libitum access to food and water. We used qPCR-based genotyping on tail biopsies to confirm hemizygous and WT genotypes for the 5xFAD strain, using the genotyping service provided by Transnetyx, Inc. (Cordova, TN, USA). All experiments were approved by the Animal Care and Use Committee (IACUC) at Houston Methodist Research Institute, Houston (Texas, USA). Studies were conducted following the NRC guide to the Care and Use of Laboratory Animals. The 5xFAD mice were randomly assigned to eight groups based on sex (male or female), injury type (Sham or TBI), and treatment (vehicle (VH) or FMT). Mice were anesthetized with isoflurane, and controlled cortical impact (CCI) injury was induced on the left hemisphere, targeting the primary motor and somatosensory cortex. This procedure was conducted using an electromagnetic Impact One stereotaxic impactor (Leica Microsystems, Buffalo Grove, IL, USA) positioned 2 mm lateral and 2 mm posterior to Bregma, with an impact depth of 1.5 mm, using a 2-mm diameter flat impact tip at a speed of 3.6 m/s and a dwell time of 100 ms, as established in our previous studies to reliably induce an inflammatory response post-TBI (Villapol et al., 2014; Villapol et al., 2017). Sham mice underwent all procedures except the impact. Mice were anesthetized and sacrificed 3 days post-injury (dpi), and brains, blood, and intestines were collected for further analysis (Figure 1a).

**Figure 1.**
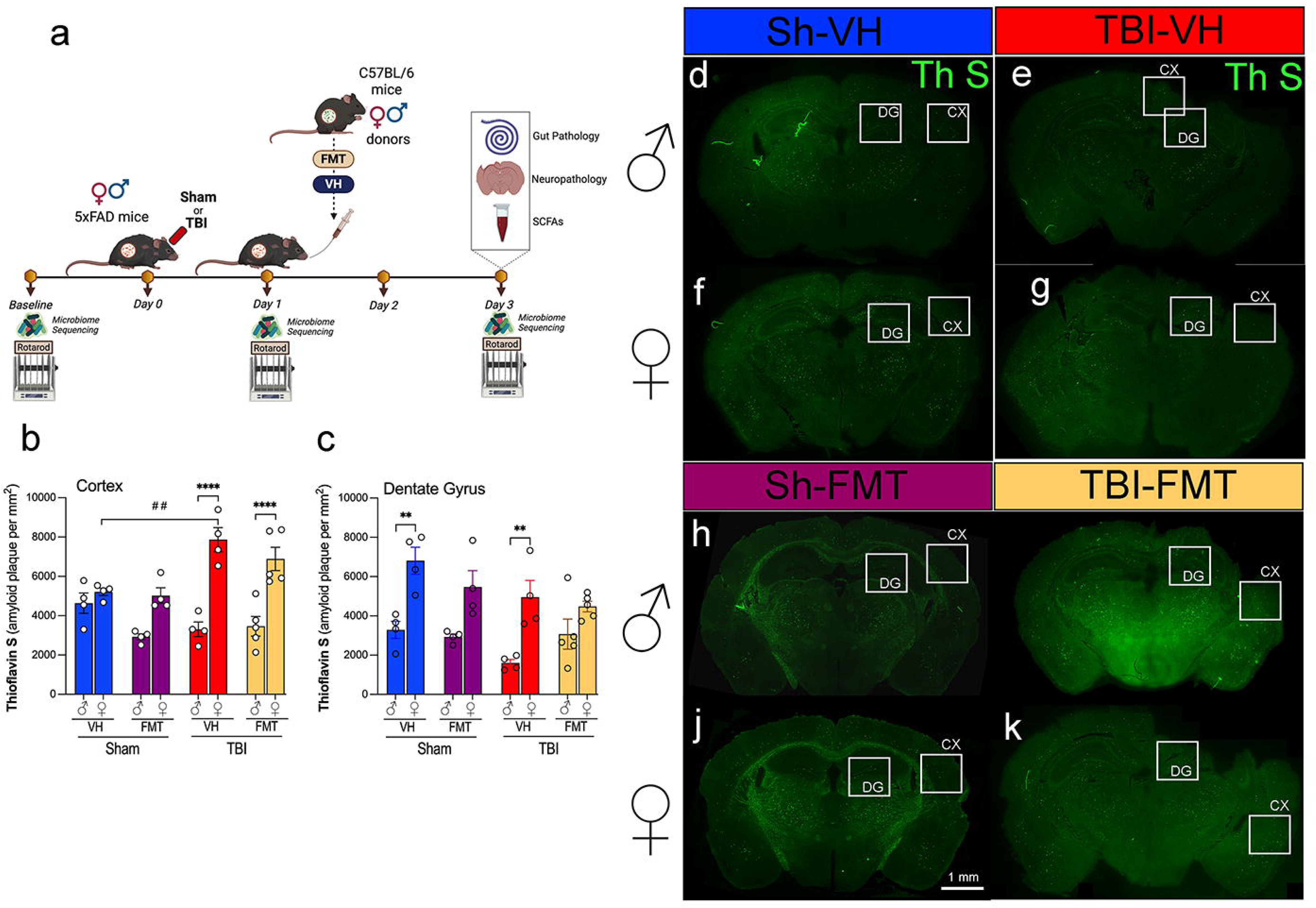
Females demonstrate greater amyloid deposition than males, with further increases in cortical regions following TBI. (a) Experimental timeline: 5xFAD male and female mice underwent baseline testing and were randomly assigned to Sham or TBI conditions. One day after injury, mice received vehicle (VH) or FMT via oral gavage from young sex-matched C57BL/6 donors. Rotarod performance and microbiome sequencing were conducted at baseline, day 1, and day 3. Brains were collected on day 3 for histological analyses. Quantification of Thioflavin-S-positive plaques in the cortex (b) and dentate gyrus (c). (d-k) Representative Thioflavin-S staining images in the cortex of male and female mice across Sham and TBI groups. Data are mean ± SEM; *p<0.05, **p<0.01, ***p<0.001, ****p<0.0001 by three-way ANOVA with Bonferroni correction for multiple comparisons. Symbols denote significant effects: *sex differences; #sham vs. TBI.

### 2.2 Rotarod Test

Sensorimotor function was evaluated using a Rotarod system (Rotamex 5, Columbus Instruments, Columbus, OH) as previously described (Villapol et al., 2012; Villapol et al., 2015). Mice received three training trials two days before testing. For each trial, mice were placed on a rotating rod for 30 s, after which the rotation speed increased from 4 to 40 rpm over 5 min. The trial ended when the mouse fell or 5 min elapsed, and the latency to fall was recorded. The mean latency from 3 trials was used as the performance measure at baseline, 1 dpi, and 3 dpi.

### 2.3 Fecal microbiota transplantation

Cecum samples were collected from 2-month-old C57BL/6 male and female mice, which served as donors for FMT. Cecal content from 3 donors of the same sex (3 males and 3 females) was homogenized in 7.5 mL of sterile phosphate-buffered saline (PBS) and centrifuged at 800 g for 3 min to pellet the large particles. The supernatants were collected in fresh sterile tubes, and the final bacterial suspension was filtered through a 70 μm strainer, collected in fresh sterile tubes, and stored at -80°C. A single 200 μL bolus of the prepared microbiota suspension was administered by oral gavage to each recipient 5xFAD mouse at 24 h after TBI (Figure 1a), by matching the sex of the FMT recipients and donors. Mice from the vehicle (VH) group received an equal volume of sterile PBS as the control group.

### 2.4 Fecal Microbiome DNA extraction

Fresh stool pellets were aseptically collected and placed in sterile tubes, immediately snap-frozen, and subsequently stored at −80°C for preservation. Genomic bacterial DNA was extracted from the frozen stool samples utilizing the QIAamp PowerFecal Pro DNA Kit (Qiagen, Germantown, MD). To facilitate DNA extraction, bead beating was conducted in three cycles, each lasting 1 min, at a speed of 6.5 m/s. There was a 5-min rest period between each cycle. This mechanical disruption was performed using a FastPrep-24 system (MP Biomedicals, Irvine, CA). Following the bead-beating process, the DNA isolation proceeded per the manufacturer’s instructions. The concentration of the extracted genomic DNA was subsequently quantified using a DS-11 Series Spectrophotometer/Fluorometer (DeNovix, Wilmington, DE).

### 2.5 Sequencing of 16S rRNA V1-V3 regions

The V1-V3 16S ribosomal RNA gene region was targeted for mouse gut microbiome characterization. The primers used for amplification contain adapters for MiSeq sequencing and single-index barcodes so that the PCR products may be pooled and sequenced directly, targeting at least 10,000 reads per sample (Caporaso et al., 2012). Primers used for the 16S V1-V3 amplification were 27F (AGAGTTTGATYMTGGCTCAG, where Y = C (90%) or T (10%); M = A (30%) or C (70%)) and 534R (ATTACCGCGGCKGCTGG, where K = G (10%) or T (90%)) (Weisburg et al., 1991). Sequencing libraries for the V1-V3 target were constructed following the instructions provided by the Illumina MiSeq system with end products of 300 bp paired-end libraries.

### 2.6 Amplicon sequence analysis pipeline

Raw data files in binary base call (BCL) format were converted and demultiplexed into FASTQs based on the single-index barcodes using the Illumina ‘bcl2fastq’ software. Forward and reverse read pairs underwent quality filtering using bbduk.sh (BBMap version 38.82), removing Illumina adapters, PhiX reads, and sequences with a Phred quality score below 15 and length below 100 bp after trimming. Quality-controlled 16S V1-V3 reads were then merged using bbmerge.sh (BBMap version 38.82), with parameters optimized for the V1-V3 amplicon (vstrict=t qtrim=t trimq=15). Further processing was performed using custom R and bash scripts, which can be found in our documentation at https://github.com/villapollab/fmt_ad_no_abx. Sequences were processed sample-wise (independent) with DADA2 v1.36 to remove PhiX contamination, trim reads (forward reads at 275 bp and reverse reads at 265 bp; reads shorter were discarded), discard reads with > 2 expected errors, correct errors, merge read pairs, and to remove PCR chimeras. After clustering, 1,839 amplicon sequencing variants (ASVs) were obtained across all samples. The ASV count table contained a total of 1,036,836 counts, at least 4,115 and at most 16,686 per sample (average 10,266). Taxonomic classification was performed in DADA2 with the assigned taxonomy function using the precompiled GreenGenes2 release 2024.09 databases (McDonald et al., 2024). The DADA2 ASV matrix, taxonomy table, and the study metadata table were combined for use within a phyloseq v1.52.0 object and merged with the phylogenetic tree externally calculated from the ASVs using MAFFT v7.525 alignment and FastTree v2.1.11 phylogenetic tree construction. The phyloseq object was used to calculate alpha and beta diversity indices, PERMANOVA calculations with vegan::adonis2 v2.6-10, relative abundance bar plots with microViz v0.12.6, R v4.5.0, and differentially abundant taxa calculations using ANCOMBC2 v2.10.1. (Lin and Peddada, 2024).

### 2.7 Serum SCFA analysis

SCFAs were analyzed at 3 dpi by a derivatization procedure. 40 µL of collected blood serum was added to 40 µL of acetonitrile, vortexed, and centrifuged. 40 µL of the supernatant, 20 µL of 200 mM 12C6-3-Nitrophenylhydrazine (3NPH), and 120 mM 1-Ethyl-3-(3-dimethyl aminopropyl) carbodiimide (EDC) were combined. 20 µL of hydrochloric acid was added and incubated for 30 min at 40°C. The resulting mixture was cooled and made up to 1.91 mL with 10% aqueous acetonitrile. 5 µL of the sample was injected into LC/MS/MS. SCFAs were separated using mobile phases: 0.1% formic acid in water (mobile phase A) and 0.1% formic acid in acetonitrile (mobile phase B). Separation of metabolites was performed on Acquity UPLC HSS T3 1.8 um (2.1×100mM). The SCFA were measured in ESI negative mode using a 6495 triple quadrupole mass spectrometer (Agilent Technologies, Santa Clara, CA) coupled to an HPLC system (Agilent Technologies, Santa Clara, CA) with multiple reaction monitoring (MRM). The acquired data was analyzed using Agilent Mass Hunter quantitative software (Agilent Technologies, Santa Clara, CA).

### 2.8 Immunofluorescence analysis

Serial free-floating coronal brain sections, 16 μm thick, were prepared at the level of the dorsal hippocampus for immunohistochemical analysis. The sections were processed using immunohistochemistry protocols, which included three consecutive 5 min washes in PBS containing 0.5% Triton X-100 (PBS-T). Sections were then treated with 5% normal goat serum (NGS) in PBS-T to block nonspecific binding for 1 h at room temperature. Overnight incubation at 4°C followed, using 3% NGS in PBS-T with primary antibodies targeting anti-rabbit Iba-1 (Wako, #019-19741, 1:500), anti-mouse Aβ6E10 (BioLegend, CAT#803001, 1:500), and anti-mouse GFAP (Millipore, CAT#MAB360, 1:500). The following day the sections were washed 3 times for 5 min each in PBS-T and incubated with the corresponding secondary antibodies (all 1:1000, Invitrogen), for 2 hours at room temperature. The sections were then rinsed with PBS three times for 5 min each and incubated in PBS with DAPI solution (1:50,000, Sigma-Aldrich, St. Louis, MO) to counterstain nuclei. The sections were rinsed with distilled water and covered with Fluoro-Gel with Tris Buffer mounting medium (Electron Microscopy Sciences, Hatfield, PA). Quantitative analysis of immunolabeled sections was performed using unbiased standardized sampling techniques across an average of three coronal sections per animal, centered on the lesion epicenter. Analyses focused on the primary somatosensory cortex and hippocampal dentate gyrus, with n=4-5 per group. Image analysis of the staining in the cortical regions was performed using ImageJ software, as previously described (Villapol et al., 2017).

### 2.9 Thioflavin-S Staining

Coronal brain sections (16 µm thick) spanning the hippocampus and cortex were mounted onto gelatin-coated glass slides (SuperFrost Plus, Thermo Fisher Scientific, IL) and air-dried. Slides then underwent a graded ethanol rehydration series (100%, 95%, 70%, and 50%) followed by two washes in distilled water (3 min each). Sections were incubated in 1% aqueous Thioflavin-S solution (Sigma-Aldrich, St. Louis, MO) for 8 minutes in the dark and subsequently differentiated by two washes in 80% ethanol and one wash in 95% ethanol (3 min each). After differentiation, sections were rinsed in three exchanges of distilled water and coverslipped with Fluoro-Gel with Tris Buffer mounting medium (Electron Microscopy Sciences, Hatfield, PA) to preserve fluorescence. Thioflavin-S-positive amyloid plaques were visualized using an epifluorescence microscope (excitation 450-490 nm, emission 515-565 nm. For quantification, images were acquired from 3-4 sections per animal at matched rostro-caudal levels and analyzed using ImageJ software. Amyloid burden was expressed as the percentage area occupied by Thioflavin-S-positive deposits within defined regions of interest (ROI).

### 2.10 Lesion volume

Brain sections were stained with Cresyl-violet. An average of 10-12 coronal brain sections, evenly spaced between 0 and −2.70 mm relative to bregma, were selected for cresyl violet staining to visualize injury-associated regions. Sections were mounted onto gelatin-coated glass slides (SuperFrost Plus, Thermo Fisher Scientific, IL) and immersed in a 1% cresyl violet solution (Sigma-Aldrich, St. Louis, MO), freshly prepared in distilled water and filtered before use. After staining, slides were sequentially dehydrated in graded ethanol solutions (50%, 70%, 95%, and 100%; 2 min each), cleared in xylene (2 × 2 min), and coverslipped using Permount mounting medium (Thermo Fisher Scientific) for long-term preservation. Lesion volume was obtained by multiplying the sum of the lesion areas by the distance between sections. The percent of lesion volume was calculated by dividing each lesion volume by the total ipsilateral hemisphere volume (similarly obtained by multiplying the sum of the areas of the ipsilateral hemispheres by the distance between sections).

### 2.11 Gut histopathological analyses

The small intestines were isolated, combined by experimental group, and fixed O/N in modified Bouin’s fixative (50% ethanol, 5% acetic acid in distilled water). Then, the ilia were prepared using the Swiss-rolling technique and kept in 4% paraformaldehyde until further processing. Tissue samples were processed using a Shandon Excelsior ES Tissue Processor and embedded in paraffin with a Shandon HistoCenter Embedding System, following the manufacturer’s standard protocols. The samples were sectioned at a thickness of 5 μm and mounted onto glass slides. Hematoxylin and Eosin (H&E) staining was performed to assess tissue structure. Intestinal sections were deparaffinized in xylene, rehydrated in water, and stained with hematoxylin for 6 hours at 60–70°C. After rinsing with tap water to remove excess stain, the sections were differentiated using 0.3% acid alcohol for 2 min, followed by eosin staining for 2 min. The slides were then rinsed and mounted with a xylene-based Permount mounting medium, allowing them to dry overnight. Alcian Blue staining was conducted to evaluate mucin production by goblet cells in the intestines. Deparaffinized sections were dehydrated in graded ethanol solutions and washed in distilled water before applying the Alcian Blue solution for 30 min. Excess stain was removed with tap water, followed by counterstaining with Nuclear Fast Red Solution for 5 min. Samples were then rinsed, dehydrated, and cleared in xylene.

### 2.11 Statistical analysis

To evaluate the combined effects of sex, injury status (sham vs. TBI), and treatment (VH vs. FMT), three-way ANOVAs were conducted with Bonferroni post hoc testing on pairwise comparisons differing by a single factor. For post-TBI comparisons, two-way ANOVAs were performed to assess the effects of sex and treatment, followed by Bonferroni’s multiple comparisons post hoc test. Non-parametric Kruskal-Wallis tests were used for microbiota diversity data. All mice were randomly assigned to experimental conditions, and experimenters were blinded to the treatment groups throughout the study. Statistical analyses were conducted using GraphPad Prism 9 software. (GraphPad, San Diego, CA, USA). Data are presented as meanL±Lstandard error of the mean (SEM), and significance levels were set at \**p*L<L0.05, \*\**p*L<L0.01, \*\*\**p*L<L0.001. Symbols in figures denote significance as follows: (*) indicates sex differences, (#) indicates differences between sham and TBI groups, and (^lZI^) denotes treatment effects in the ANOVAs post-hoc tests with Bonferroni correction.

## 3. Results

### 3.1 Sex-Specific effects on amyloid deposition in the cortex and dentate gyrus post-TBI

To investigate the impact of FMT on amyloid pathology following TBI, we used 5xFAD mice subjected to sham or TBI surgery and subsequently transplanted with VH or microbiota from C57BL/6 donor mice (Figure 1a). Microbiome sequencing and motor testing were conducted at baseline, 1 dpi and 3 dpi, and tissue analysis was performed at endpoint (3 dpi).

Thioflavin-S (Thio-S) staining revealed abundant amyloid plaque deposition in both the cortex and dentate gyrus (DG) of 5xFAD mice (Figure 1b-e). Quantification showed that cortical plaque burden was significantly increased in female TBI mice compared to female sham controls (##p<0.01), exhibiting higher amyloid burden after injury (Figure 1f). At 3 dpi, female TBI mice exhibited a significantly greater cortical amyloid plaque burden compared to their male counterparts treated with either VH or FMT (****p < 0.0001). FMT did not alter the cortical amyloid burden substantially relative to VH controls. In the DG, amyloid plaque burden was similarly elevated in female VH mice compared to male VH controls under both sham and TBI conditions (Figure 1g). However, this difference was not seen in the FMT groups. Together, these data indicate that the amyloid accumulation in 5xFAD mice occurs in a region- and sex-specific manner, and that TBI synergizes with microbial transfer to exacerbate cortical plaque pathology.

### 3.2 FMT modulates A**β** pathology in a sex- and brain region-specific manner

To further evaluate how FMT influences Aβ pathology following TBI, we performed immunostaining with the 6E10 antibody in cortical and hippocampal regions of 5xFAD mice (Figure 2a-d). In sham animals, both male and female mice showed comparable levels of cortical and DG immunoreactivity, with no significant differences between VH and FMT groups (Figure 2a, b, e). Quantitative analysis of 6E10 immunoreactivity revealed a significant increase in cortical plaque load in both TBI males (####p < 0.0001 and ##p < 0.01 for VH and FMT groups, respectively) and females (###p < 0.001 for FMT group) compared to their sham counterparts, with a greater increase in TBI-VH males compared to females (*p < 0.05) (Figure 2e). Moreover, Figure 2e also shows that FMT reduced the 6E10 immunoreactivity in VH-treated TBI-male mice (^lZI^p< 0.05). In the DG, both male and female TBI groups significantly displayed elevated 6E10+ area relative to sham controls, (Figure 2f). FMT did not significantly alter amyloid levels in sham or TBI groups. The expression of 6E10 shows in the form (Figure 2a1-d4) of Aβ deposits with plaque morphology. To further characterize plaque maturation, Aβ deposits were classified into three morphologies (Figure 2g): diffuse plaques (large, ill-defined), compact plaques (sharply delineated), and dense-core plaques (fibrillar, with a compact core and surrounding halo) (Thal et al., 2002; Serrano-Pozo et al., 2011; Perez-Nievas et al., 2013; Viejo et al., 2022). Dense-core plaques have been associated with neuritic dystrophy and gliosis, reflecting advanced AD pathology (Serrano-Pozo et al., 2011). In the cortex, TBI induced a marked decrease in total plaque numbers in male mice compared to Sham controls (#p < 0.05 for VH and FMT groups) (Figure 2h). Subtype analysis further revealed that dense-core plaques were significantly reduced in Sham males treated with FMT compared to VH (^lZI^p < 0.05) (Figure 2h). Similarly, Sham females receiving FMT showed decreases in diffuse and dense-core Aβ burden relative to VH controls (^lZI^p < 0.05, respectively), whereas no treatment differences were observed in the TBI groups (Figure 2j). In the DG, morphological analysis indicated that in females, significant differences were observed in the diffuse subtype, with a reduction in Sham FMT compared to Sham VH mice (^lZI^p < 0.05) (Figure 2k) that was absent in TBI groups and in males (Figure 2i).

**Figure 2.**
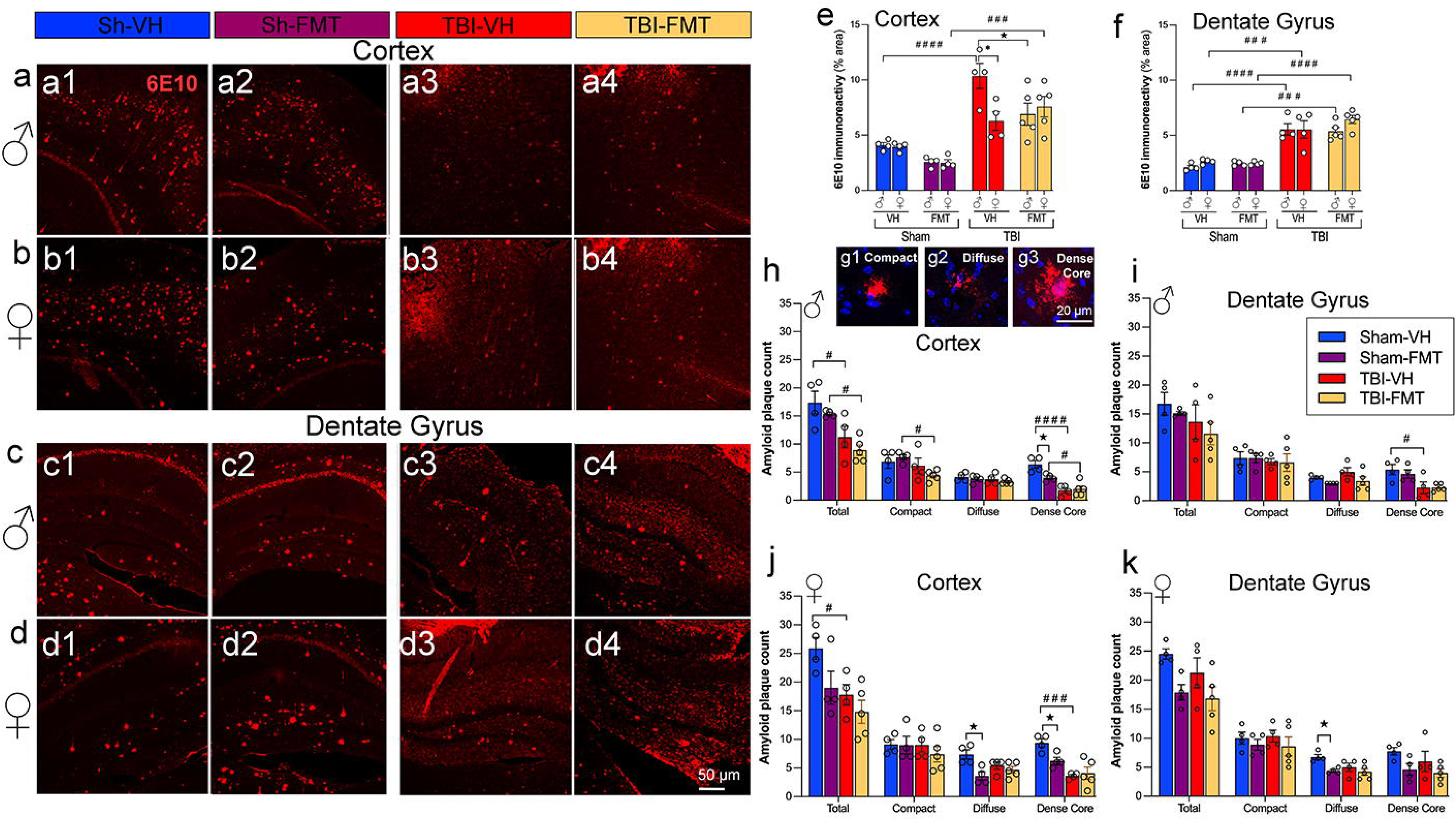
FMT modulates Aβ plaque burden and morphology after TBI. (a-d) Representative 6E10 immunostaining of cortical and dentate gyrus sections from male and female 5xFAD mice across Sham-VH, Sham-FMT, TBI-VH, and TBI-FMT groups. (e) Quantification of plaque intensity revealed significant increases after TBI in both sexes, with FMT reducing plaque levels in males in the cortex. Notably, 6E10 cortical staining showed a significant reduction between male TBI-FMT and male TBI-VH. (g1-3) Plaque morphology was categorized into compact, diffuse, and dense core types. (h, j) In the cortex of males, dense-core plaque burden was significantly reduced between Sham-VH and Sham-FMT groups in cortex and DG. (j, k) In females, diffuse plaque burden differed significantly between VH-Sham and FMT-Sham groups (p < 0.01). Overall, TBI shifted plaques toward a more diffuse phenotype with a concomitant reduction in dense-core plaques, an effect observed across both VH- and FMT-treated mice in males and females. Data are presented as mean ± SEM. *p < 0.05, **p < 0.01, ***p < 0.001, ****p < 0.0001 by three-way ANOVA (6E10 immunoreactivity) or two-way ANOVA (plaque density, by sex and plaque type) with Bonferroni correction. Symbols indicate significance: *sex differences, #sham vs. TBI, and ^lZI^treatment effects.

Taken together, these findings demonstrate that injured brains exhibit reduced amyloid pathology in 5xFAD mice in a sex-dependent manner, with females showing greater susceptibility. Importantly, the effects of FMT were most apparent in Sham AD mice, where amyloid deposition was reduced. In contrast, in TBI animals, the neuroinflammatory and neurodegenerative environment appeared to override potential microbiome-mediated benefits.

### 3.3 FMT reduces glial activation in the cortex of 5xFAD male mice following TBI

To evaluate the effect of FMT on neuroinflammation after injury, we quantified Iba-1 and GFAP immunoreactivity in the cortex and DG of 5xFAD mice. Following TBI, male mice treated with FMT showed decreased Iba-1+ microglial activation in the cortex and DG compared to VH controls. However, FMT treatment had no effect in female mice (Figure 3a, b). GFAP+ astrocytic reactivity was decreased in male TBI mice treated with FMT compared to females in the cortex, while no changes were observed in any groups for the DG (Figure 3c, d). These findings indicate that FMT modulates microglial and astrocytic responses after TBI in a sex- and region-specific manner.

**Figure 3.**
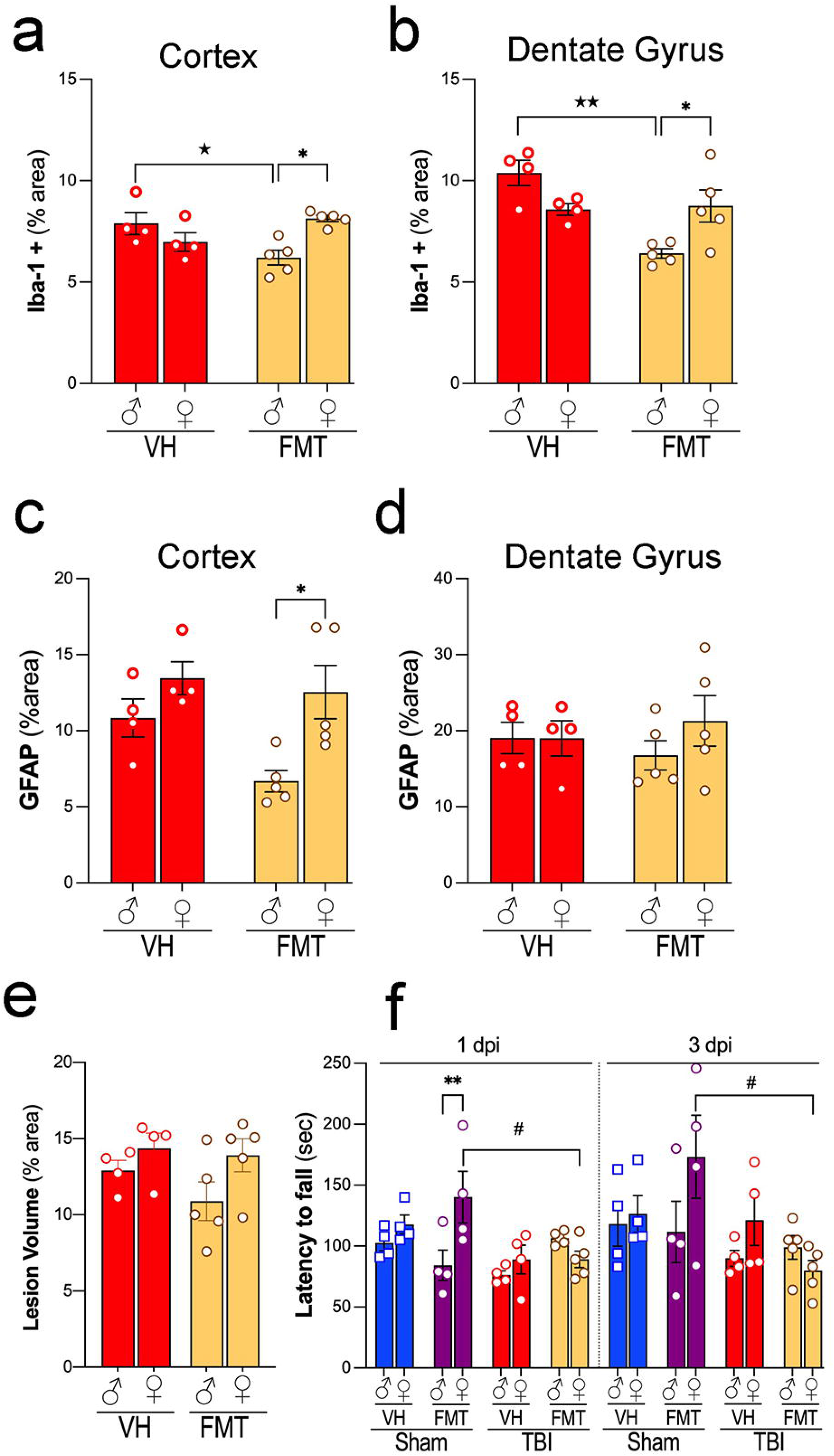
Sex-differences in the impact of FMT on neuroinflammation, lesion volume, and motor outcomes after TBI in AD mice. (a–b) Quantification of Iba1+ microglia in the cortex (a) and dentate gyrus (b) shows increased microglial activation in the VH compared to the FMT group in the male mice at 3 dpi (^lZI^p < 0.05, ^lZIlZI^p < 0.01). (c–d) GFAP+ astrocyte reactivity in cortex (c) and dentate gyrus (d) demonstrates an increase in the females versus males only in the FMT group and not differences between the treatments at 3 dpi. (e) Lesion volume analysis reveals no significant differences between VH and FMT groups for both sexes. (f) Motor performance assessed by rotarod (latency to fall) at 1 dpi and 3 dpi shows sex-dependent impairment after TBI, with reduced latency observed only in FMT-treated females (#p < 0.05 and **p< 0.01, respectively). Data are presented as mean ± SEM. Statistical significance was determined by two-way (Iba-1 and GFAP % area, and lesion volume) or three-way (latency to fall in the rotarod) ANOVA with Bonferroni correction for multiple comparisons. Symbols denote significant effects: *sex differences; # sham vs. TBI, and ^lZI^treatment effects.

Next, we assessed whether FMT influenced structural injury and functional outcomes. Lesion volume, measured as percent damaged cortical area, did not differ significantly between VH- and FMT-treated TBI groups in either sex (Figure 3e). However, motor performance on the rotarod revealed that Sham females treated with FMT exhibited improved latency to fall compared to VH at 1 dpi whereas in TBI groups, FMT treatment was associated with reduced performance at both 1 and 3 dpi compared to VH controls (#p < 0.05) (Figure 3f). Moreover, Sham-FMT female mice performed significantly in the rotarod test compared to Sham-FMT male mice at 1 dpi (**p < 0.01).

### 3.4 FMT modifies gut microbial community profiles after TBI without altering alpha diversity or richness

To investigate the impact of TBI and FMT on gut microbiota, we first assessed alpha diversity and richness across experimental groups. We investigated the effects of FMT on gut microbial diversity and composition in 5xFAD mice subjected to sham or TBI procedures. Measures of alpha diversity (Shannon index) revealed no significant differences between groups in either males (Figure 4a) or females (Figure 4b) at 1 or 3 dpi. Similarly, microbial richness was not significantly altered across experimental groups at either time point (Figure 4c–d), with or without FMT treatment. We next evaluated beta diversity using principal coordinate analysis (PCoA) based on Bray–Curtis dissimilarity (Figure 4e-g), showing marked group-dependent effects. At baseline, microbial communities did not differ significantly among groups (Figure 4e; Pr(>F) = 0.207). However, by 1 dpi, significant separation was observed across sham and TBI groups (Figure 4f; Pr(>F) = 0.003, R² = 0.228), with clustering driven by both injury and FMT status. This effect was more pronounced at 3 dpi (Figure 4g; Pr(>F) = 0.001, R² = 0.514), where distinct microbiome profiles were evident between sham and TBI mice, and between VH and FMT-treated animals, indicating that donor microbiota exerted a measurable effect on post-injury microbial engraftment.

**Figure 4.**
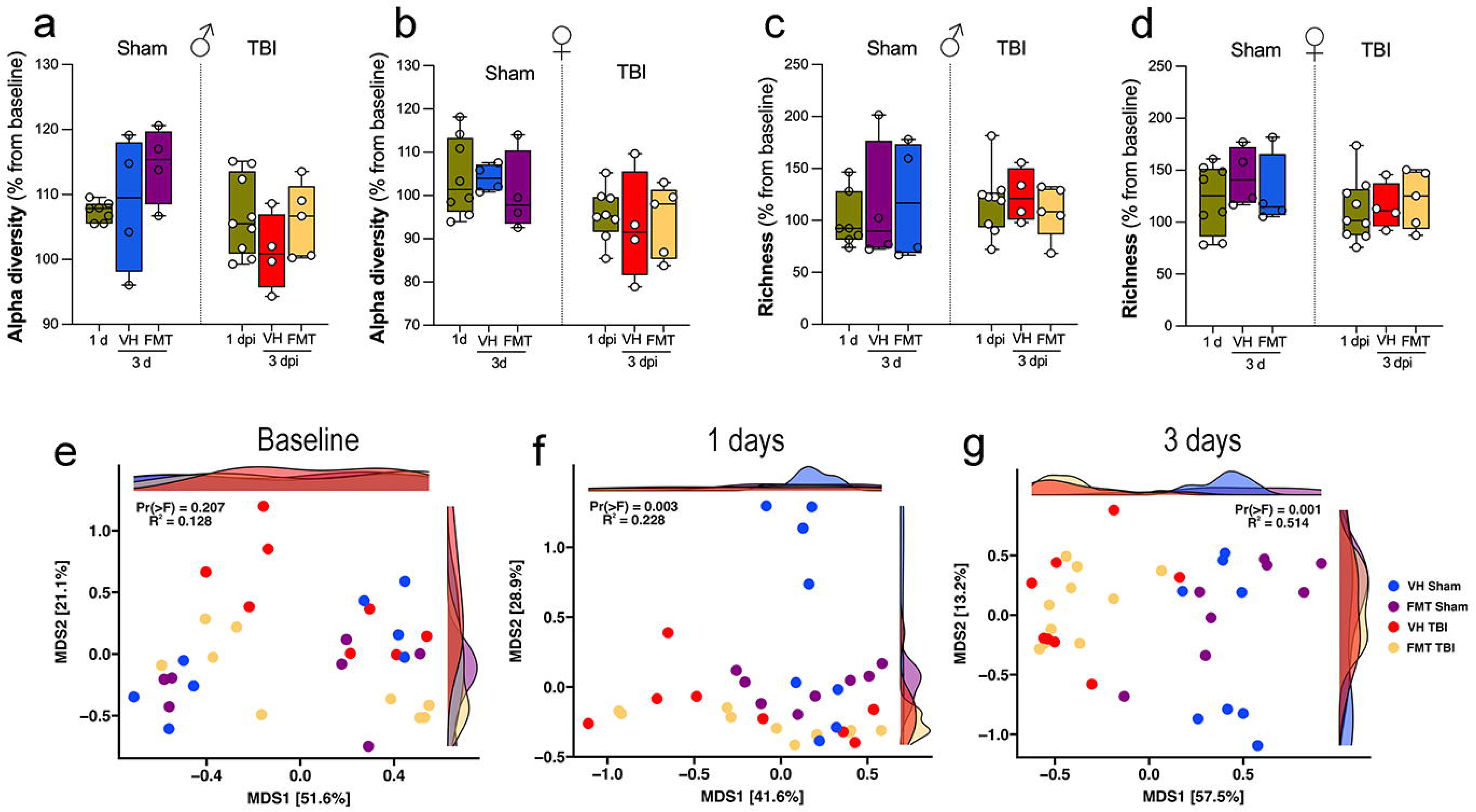
FMT modulates gut microbiome diversity and composition after TBI in a sex-dependent manner. Difference in diversity and richness of fecal microbiota compared to baseline at 1 day and 3 days post-intervention. No significant differences were detected in diversity (a, b; Shannon index) or richness (c, d; Chao1 index) across sham and TBI groups, stratified by sex and treatment (VH vs. FMT). Data are presented as mean ± SEM. Alpha diversity and richness were analyzed with Kruskal-Wallis. (e-g) Beta diversity analysis of gut microbial communities by principal coordinate analysis (PCoA) based on Bray-Curtis distances at baseline (e), 1 day (f), and 3 days (g) post-intervention. Each point represents an individual sample, colored by group (sham-VH, sham-FMT, TBI-VH, TBI-FMT) and sex. Density plots show the distribution along MDS1 and MDS2 axes. At 3 dpi, a significant separation was seen between the microbial communities of the VH TBI and FMT TBI mice compared to VH-sham and FMT sham mice (PERMANOVA).

Together, these results indicate that while FMT and TBI do not significantly alter alpha diversity or richness, they strongly reshape the overall microbial community structure, with effects becoming increasingly evident over time after injury. FMT partially modulates these shifts, suggesting that microbiota restoration strategies may counteract aspects of TBI-induced dysbiosis in a sex-dependent manner.

### 3.5 FMT modulates microbial taxonomic composition dynamics following TBI

To investigate specific microbial shifts associated with FMT after TBI, we analyzed genus-level relative abundances across groups and time points. Heatmap clustering revealed distinct temporal patterns in Sham VH, TBI VH, Sham FMT, and TBI FMT mice (Figure 5a-d). Sham VH mice showed relatively stable microbial profiles over time, with moderate fluctuations in genera such as *Lepageella*, *Pelethenecus*, *Muribaculaceae* family members, *Oscillospiraceae*, and *Acetatifactor*, were consistently detected across timepoints (Figure 5a, e). In contrast, TBI VH mice exhibited marked temporal changes, with increased abundance of *Duncaniella*, *Bacteroides_H_857956*, and *Ligilactobacillus* by 3 dpi (Figure 5b) and with a decrease in beneficial taxa such as *Lactobacillus*, *Turicibacter*, and *Dubosiella* (Figure 5b, d).

**Figure 5.**
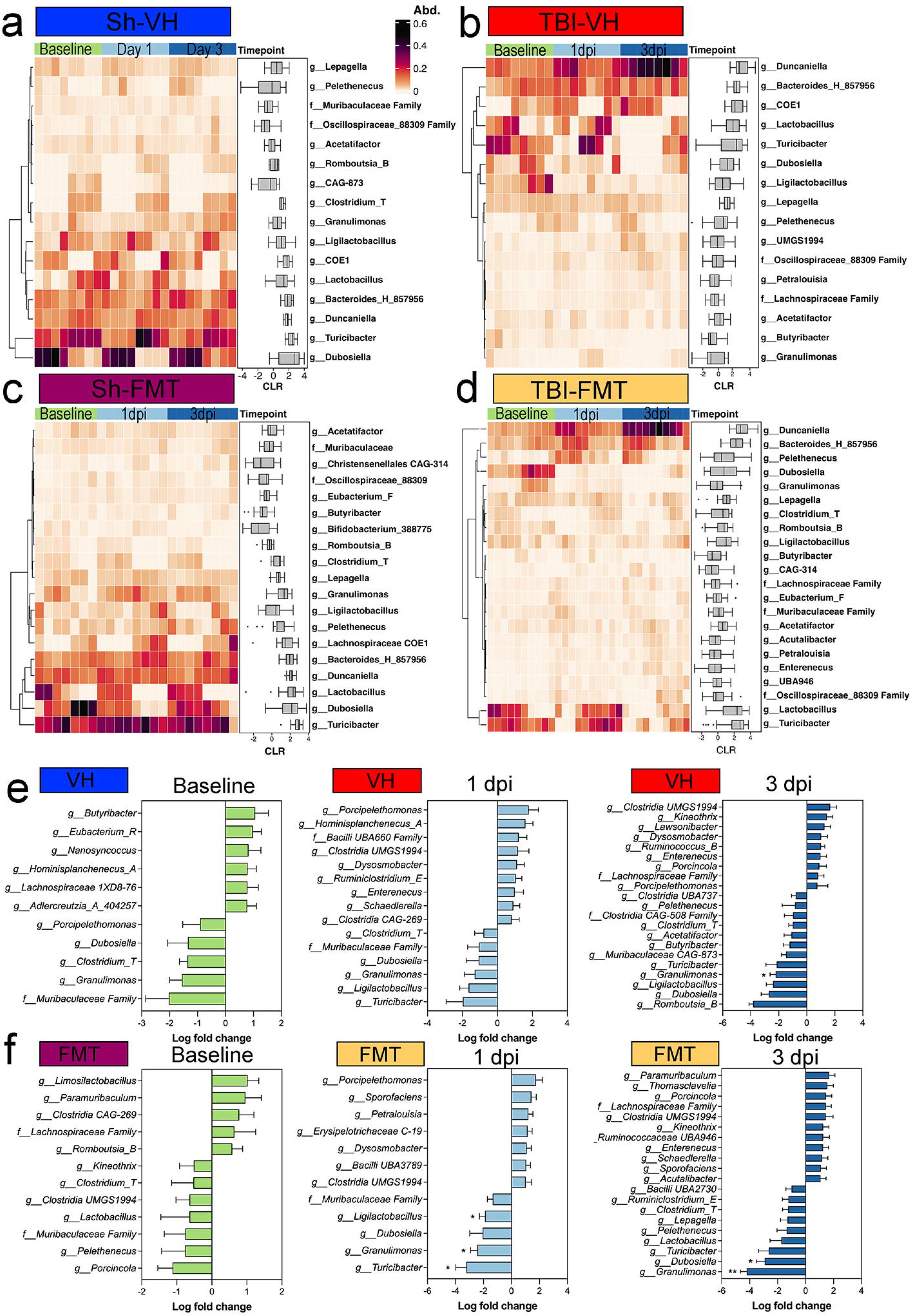
Taxonomic changes in fecal microbiome composition in 5xFAD mice following TBI and FMT. Heatmaps showing the relative abundance (Abd.) of bacterial genera at baseline, 1 day, and 3 days post-intervention in (a) Sham-VH, (b) TBI-VH, (c) Sham-FMT, and (d) TBI-FMT groups. Color scale represents relative abundance (0-0.6). Boxplots adjacent to each heatmap display centered log-ratio (CLR) transformed abundance changes over time for key taxa. Differential abundance analysis of Sham and TBI mice within each treatment group at each timepoint identified taxa significantly altered within each group, shown as log fold changes for (e) VH treatment and (f) FMT. Negative values indicate decreased abundance relative to Sham; positive values indicate enrichment. In TBI-FMT mice, reductions were observed in *Turicibacter, Granulimonas,* and *Ligilactobacillus* at the 1dpi timepoint, with the altered *Granulimonas* persisting into the 3 dpi timepoint of both VH and FMT groups.

FMT treatment reshaped microbial trajectories in both sham and injured animals. Sham FMT mice demonstrated enrichment of beneficial taxa, including *Bifidobacterium*, *Butyricicoccus*, and *Christensenellaceae* family members, particularly by 3 dpi (Figure 5c). In TBI FMT mice, the microbial community was dominated by reduction in SCFA-producing taxa, including *Turicibacter*, *Granulimonas*, and *Ligilactobacillus*, when comparing Sham and TBI cohorts (Figure 5e, f).

FMT treatment produced distinct patterns of microbial remodeling depending on the condition. At 1 dpi, VH mice exhibited only minor, non-significant changes in bacterial genera. In contrast, FMT-treated mice showed a significant reduction in SCFA-producing taxa, including *Turicibacter*, *Granulimonas*, and *Ligilactobacillus*, when comparing Sham and TBI cohorts (Figure 5e, f). By 3 dpi, differences between Sham and TBI groups persisted: *Granulimonas* remained reduced, and another key gut–brain axis taxon, *Dubosiella*, was significantly depleted in FMT-treated TBI animals. Together, these findings indicate that TBI disrupts gut microbial composition by selectively enriching pro-inflammatory species while depleting protective commensals. FMT partially counteracts these effects by supporting restoration of beneficial taxa, particularly SCFA producers such as *Dubosiella*. These taxonomic shifts suggest potential mechanisms by which microbiota modulation may influence host neuroinflammatory outcomes following TBI.

### 3.5 TBI or FMT did not significantly alter SCFA levels

We next examined how FMT affected serum SCFA composition in sham and TBI 5xFAD mice. Among the measured SCFAs, isovalerate levels were significantly elevated in Sham-FMT in females compared to male mice (Figure 6a; **p < 0.01). Other SCFAs, including valerate (Figure 6b), caproate (Figure 6c), isobutyrate (Figure 6d), butyrate (Figure 6e), propionate (Figure 6f), 2-methylbutyrate (Figure 6g), 3-methylvalerate (Figure 6h), and isocaproate (Figure 6i) remained unchanged across treatments, sexes and injury conditions.

**Figure 6.**
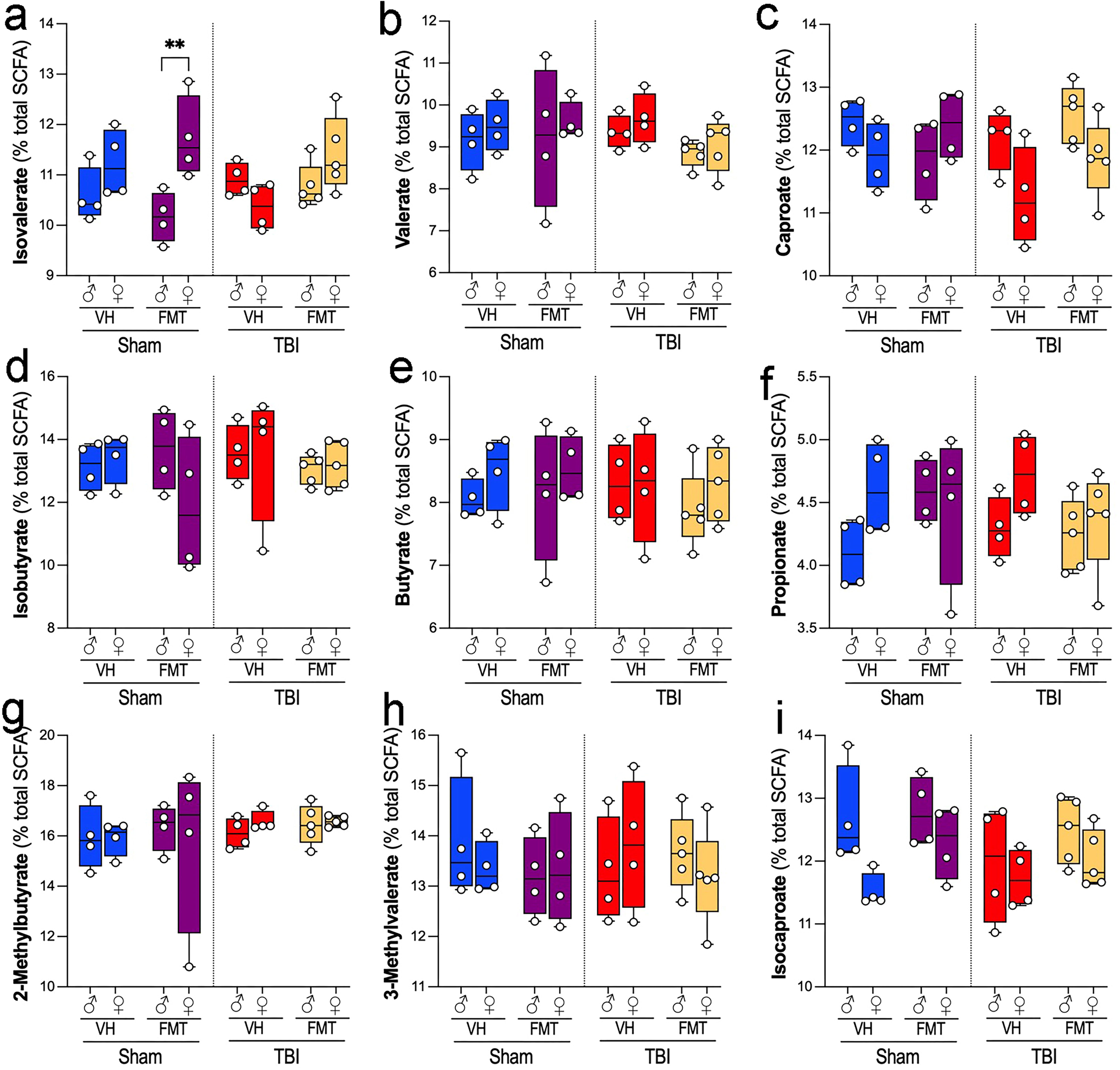
Effects of TBI and FMT on short-chain fatty acid (SCFA) profiles in male and female 5xFAD mice. Relative abundance of fecal SCFAs (% of total SCFA pool) was quantified in male and female 5xFAD mice subjected to sham or TBI surgery and treated with VH or FMT. Panels show: (a) isovalerate, (b) valerate, (c) caproate, (d) isobutyrate, (e) butyrate, (f) propionate, (g) 2-methylbutyrate, (h) 3-methylvalerate, and (i) isocaproate. Sham FMT females had a higher abundance in isovalerate (b; **p < 0.01) compared to sham FMT males, while no significant differences were observed in the TBI groups. Other SCFAs remained unchanged across treatment and injury conditions. Data are presented as mean ± SEM. Statistical significance was determined by three-way ANOVA with Bonferroni correction for multiple comparisons. Symbols denote significant effects: *sex differences and #sham vs. TBI.

### 3.6 FMT modulates gut morphology in 5xFAD male mice

To investigate whether TBI and FMT influence gut structural integrity, we analyzed histological parameters of the small intestine and colon using H&E-stained sections (Figure 7a-h). Representative H&E images revealed preserved villus and crypt structures across all groups, although subtle alterations were evident between sham and TBI conditions.

**Figure 7.**
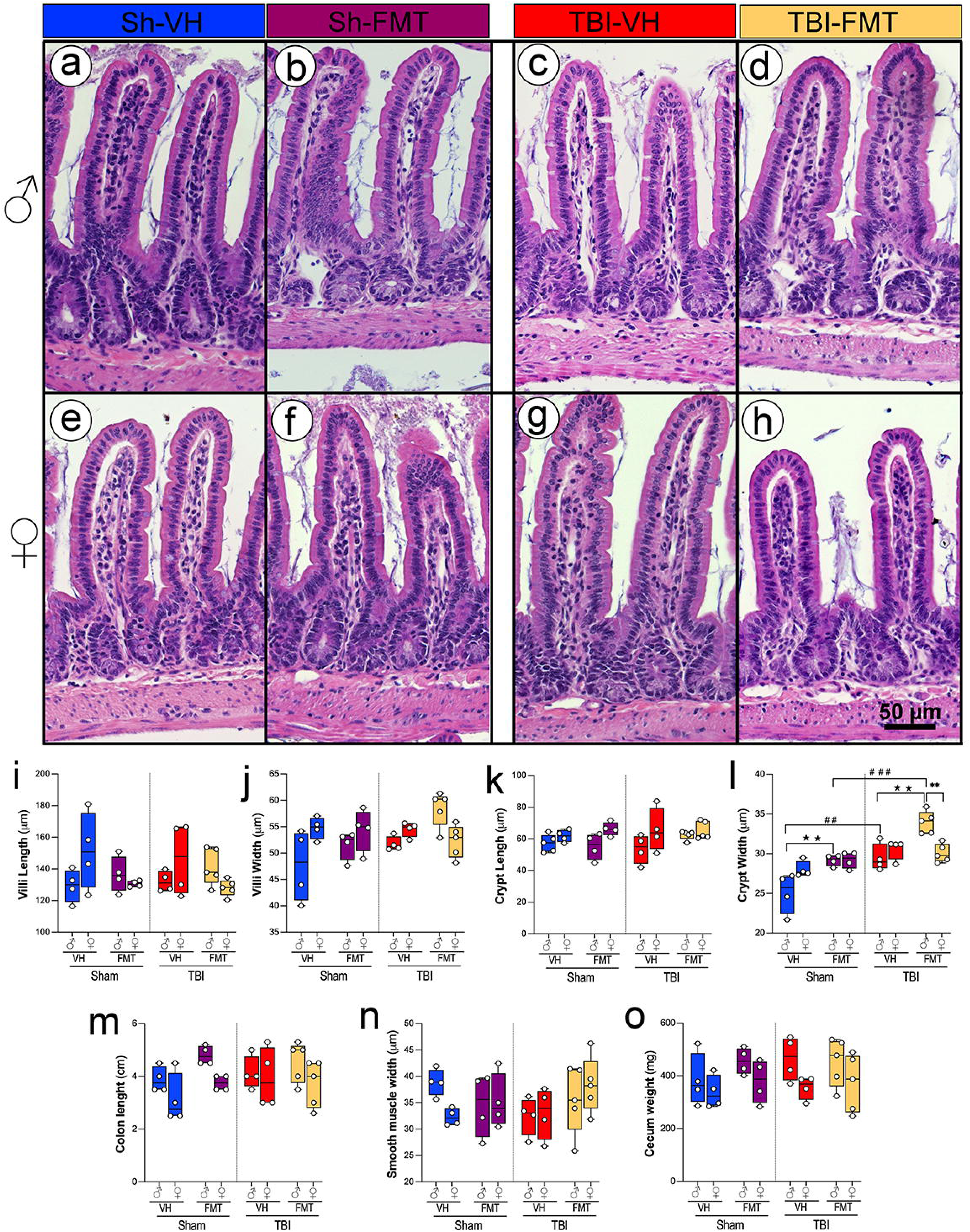
Effects of TBI and FMT on intestinal morphology in male and female 5xFAD mice. Representative H&E images of small intestinal villi and crypts from male (a-d) and female (e-h) 5xFAD mice across groups: Sham VH (a, e), Sham FMT (b, f), TBI VH (c, g), and TBI FMT (d, h). Quantification of villus length (i), villus width (j), crypt length (k), and crypt width (l) revealed no differences in villus or crypt length and villus width between groups. However, crypt width was significantly increased in TBI mice compared to Sham controls, particularly in VH- and FMT-treated males (##p < 0.01, ###p < 0.001). Gross gut anatomy measures, including colon length (m), smooth muscle thickness (n), and cecum weight (o), were not significantly altered by either TBI or FMT. Data are shown as box-and-whisker plots with individual values. Symbols denote significant effects: *sex differences, #Sham vs. TBI, and IZtreatment effects.

Quantitative analysis showed no significant differences in villus length or villus width across groups (Figure 7i-j). Similarly, crypt length remained unchanged in both sham and TBI animals regardless of treatment (Figure 7k). In contrast, crypt width was significantly increased in TBI mice compared to sham controls, particularly in males with both VH and FMT treatments (Figure 7l; ##p < 0.01, ###p < 0.001). FMT treatment attenuated this enlargement, increasing crypt width only in males in both Sham and TBI groups (IZIZp < 0.01). In addition, in the TBI group, an increase is observed in the males compared to females only in the FMT-treated mice, suggesting that FMT modifies epithelial architecture dependent on injury (**p<0.01).

Other gross measures of gut anatomy, including colon length, smooth muscle thickness, and cecum weight, did not differ significantly between experimental groups (Figure 7m-o).

Together, these results demonstrate that while overall gut architecture remains intact following TBI, crypt morphology is selectively altered, and FMT modulates these changes, suggesting a sex-specific susceptibility and protective role in maintaining epithelial structure after injury.

## 4. Discussion

In this study, we subjected AD mouse models to TBI and found that the injury accelerated amyloid pathology, particularly in females. A short-term, single FMT administration was not sufficient to counteract the TBI-induced neuroinflammatory response, although notable effects on the amyloid pathology were observed in sham AD mice. These findings indicate that a single FMT intervention cannot fully reverse TBI-associated neuroinflammation, but it does elicit sex-dependent alterations in brain amyloid pathology and microbiome profiles.

### 4.1 TBI exacerbates amyloid pathology and neuroinflammation in a sex-dependent manner

Consistent with previous reports linking TBI to accelerated AD progression (Ramos-Cejudo et al., 2018), our results have shown an increase in cortical amyloid accumulation, increased Aβ plaques, and impaired motor performance. These effects were more pronounced in females, reinforcing clinical and preclinical evidence of heightened vulnerability to AD pathology in females (Zhu et al., 2021; Vila-Castelar et al., 2025).

Our findings demonstrate that TBI significantly increased overall Aβ6E10 immunoreactivity in the cortex and hippocampus of both male and female mice. Notably this increase in total amyloid burden was accompanied by a relative reduction in compact and dense-core plaques. Given that soluble Aβ oligomers are widely regarded as the most neurotoxic species, whereas dense-core fibrillar plaques may serve to sequester soluble forms, this morphological shift suggests altered aggregation dynamics rather than a protective effect (Goure et al., 2014; Xu et al., 2020). Rather than mitigating pathology, the reduction of dense-core plaques after TBI may indicate impaired sequestration of soluble Aβ, thereby favoring accumulation of oligomeric and diffusely aggregated forms that are more harmful to AD pathology. Thus, TBI appears to exacerbate amyloid pathology through enhanced accumulation of aggregated Aβ, implicating secondary injury mechanisms in the worsening of neurodegenerative processes.

### 4.2 Impact of FMT on the neuropathology post-TBI

While prior studies reported that FMT-treated mice exhibited significant preservation of cortical volume and white matter connectivity at chronic time points, such as 60 dpi (Davis et al., 2022a) or 90 dpi (Du et al., 2021), our findings did not reveal comparable effects during the acute phase. However, the interpretation of the mouse study warrants cautions due to limited statistical power. In a comparable pig model, FMT modestly enhanced recovery rates as reflected by gross MRI pathology markers and motor function (Fagan et al., 2023). Beyond their therapeutic potential, FMT has also been employed as an investigative tool to study disease mechanisms and pathology rather than solely as an intervention for recovery.

In this study, the absence of any effect on lesion size reinforces the idea that the benefits of FMT arise not from direct neuroprotection against acute mechanical trauma, but rather from modulation of secondary injury cascades. Consistent with this, our findings indicate that microbiome modulation may influence the qualitative features of amyloid pathology, such as plaque morphology, rather than entirely preventing the trajectory of Aβ deposition. This effect warrants further mechanistic investigation. TBI induced robust microgliosis and astrogliosis in both the cortex and dentate gyrus. Yet, FMT exerted significant effects, selectively attenuating microglial activity in the cortex and DG in male mice while leaving astrocytic reactivity largely unchanged. Despite these histological changes, FMT did not reduce lesion volume or improve motor outcomes, suggesting that microbiota-based interventions alone may be insufficient to alter gross injury outcomes in this aggressive AD model. Although females also exhibited greater TBI-induced amyloid pathology, they also showed unique microglial remodeling in response to FMT. These findings parallel established sex differences in immune responses (Villapol et al., 2017), underscoring the importance of developing sex-specific microbiome-based therapeutic strategies.

### 4.3 TBI disrupts microbiota beta diversity

TBI has been shown to reduce microbial diversity and richness, consistent with post-injury dysbiosis observed in both human and rodent studies (Treangen et al., 2018; Nicholson et al., 2019). In our AD model, however, we did not detect such differences following TBI, reflecting variability among samples and the limited number of mice analyzed. Analysis of microbial communities revealed that TBI altered beta diversity in a time-dependent manner. Differences emerged as early as 1 dpi and became more pronounced by day 3, with apparent clustering of TBI and sham communities. These shifts were also sex-dependent, with females showing more pronounced reductions in richness following TBI (Figure 4e-g). Community separation, evident at 1- and 3-dpi, suggests rapid and sustained ecological disruption driven by systemic inflammation, altered gut motility, and stress-related hormonal changes (El Baassiri et al., 2024).

### 4.4 16S rRNA profiling identified distinct taxonomic signatures associated with TBI and FMT, with no differences in SCFA production

FMT from healthy donors reshaped the gut microbial community, but its effects were highly context dependent. Several studies demonstrate that FMT can ameliorate TBI-induced neuropathology, attenuating lesion size, ventriculomegaly, and white matter loss while reducing microglial inflammatory gene expression (Du et al., 2021; Davis et al., 2022b; a). Translational studies extend these findings to large animal models, where FMT limited neuronal loss and improved motor outcomes in pediatric piglets (Fagan et al., 2023), highlighting its clinical potential. This sex-specific divergence highlights underlying differences in microbiome stability and responsiveness, which may contribute to distinct neuroinflammatory and pathological outcomes between males and females. In this study, there were no significantly altered taxa between the sham and TBI groups in either the VH or FMT animals at baseline. By 1 dpi, however, the TBI FMT group showed a significant decrease in *Ligilactobacillus*, *Granulimonas*, and *Turicibacter,* whereas no such significant taxonomic alterations were seen in the sham-TBI comparison of the VH group. At 3 dpi, both VH and FMT TBI mice presented with significantly decreased *Granulimonas*, with the FMT TBI mice showing significantly decreased *Dubosiella*. The reduction in both *Granulimonas* and *Dubosiella* suggests that the altered metabolic environment of the injured gut may create distinct ecological niches that favor different donor-derived microbes (Everard et al., 2013; O’Callaghan and van Sinderen, 2016). These divergent outcomes emphasize that the host context influences colonization dynamics, which may explain why FMT can be beneficial in some settings but neutral or even detrimental in others. Functional analyses of microbial metabolites provided additional insights. SCFAs, including acetate, propionate, and butyrate, are critical regulators of epithelial barrier integrity, microglial maturation, and neuroinflammatory tone (Erny et al., 2015). SCFA levels, particularly butyrate and isobutyrate, correlated with improved motor function and reduced neuroinflammation, supporting their role as neuroprotective metabolites (Dalile et al., 2019). However, we did not observed changes in the SCFA levels to sham values or treatment. These patterns can be disrupted by TBI-induced anorexia, impaired gut motility, and accelerated mucosal turnover (Opeyemi et al., 2021). Female sham-FMT animals only showed higher isovalerate compared with males, suggesting sex-specific metabolic outputs. These patterns suggest a potential functional consequence of microbiota shifts that may require larger cohorts or more extended observation periods to confirm.

### 4.5 TBI and FMT reshape intestinal morphology

TBI-induced neuropathological changes occurred alongside intestinal structural alterations, villus atrophy, crypt hypertrophy, and increased crypt width, supporting prior findings that brain injury rapidly disrupts gut homeostasis (Opeyemi et al., 2021). Our results demonstrate that TBI increases crypt width in males, but not in females, an indicator often associated with epithelial hyperplasia in response to injury or inflammation (Hang et al., 2003). This increase was evident in the sham male mice and was further exacerbated by TBI. FMT amplified some of these changes, particularly increasing villus width in TBI mice. While such structural alterations could represent adaptive remodeling to enhance absorptive surface area, they might also reflect a pro-inflammatory mucosal state if accompanied by immune cell infiltration (Luissint et al., 2016). These findings suggest that microbiota transfer affects epithelial remodeling processes, potentially through altered microbial signaling or metabolite production. TBI may further weaken morphological function for proper immune integration, which is potentially restored by FMT and results in drastic epithelial adjustments. The significant increase in crypt width after individual FMT and combined FMT-TBI exposure suggests that FMT treatment may induce short-term intestinal inflammation and epithelial remodeling, perhaps through microbial competition to re-establish a heathy gut microbiome (Pu et al., 2024). Interestingly, colon length, smooth muscle thickness, and cecum weight remained unchanged, indicating that changes were localized to epithelial than global gut anatomy. Restoration of gut structure, particularly normalization of villus and crypt architecture, may reduce systemic inflammation and microbial translocation, thereby dampening neuroimmune activation.

### 4.6 Implications and future directions

A key limitation of this study is the short post-treatment period after TBI, which captures acute but not chronic changes in the microbiome, metabolites, and neuropathology. Longitudinal studies will be essential to determine whether the effects of early FMT persist or evolve. Expanded metabolomic profiling, including bile acids, amino acid derivatives, and neurotransmitter precursors, could provide a more complete functional landscape of gut microbial activity post-TBI.

Another limitation is its relatively small sample size. Pooling mice by sex, while necessary for statistical power, may reduce sensitivity to detect subtler but biologically meaningful differences within highly interconnected systems such as the gut microbiome.

Mechanistic studies using germ-free mice or microbiota depletion before FMT treatment could help clarify causal pathways linking microbiota modulation to amyloid pathology. Given that FMT outcomes are donor-dependent, future work should aim to identify specific microbial strains or metabolites responsible for neuroprotection, enabling more targeted and standardized therapies. At the microbial species level, transplantation of *Prevotella copri* was shown to restore gut homeostasis and protect against oxidative stress (Gu et al., 2024), further supporting the therapeutic capacity of targeted single microbial interventions. Complementary metabolomic approaches identified natural compounds derived from gut microbial metabolism as potent inhibitors of TBI-induced microglial NLRP3 inflammasome activation, which should be interesting to apply (Tang et al., 2025).

## 5 Conclusion

Collectively, we have shown that TBI accelerates amyloid pathology, neuroinflammation, and gut dysbiosis in the 5xFAD model, with females displaying greater vulnerability. FMT partially enriched beneficial taxa and altered plaque morphology but failed to prevent amyloid accumulation, microgliosis, or behavioral deficits. These results underscore the gut–brain axis as a key link between TBI and AD and highlight the need for longer-term, combined, and sex-specific microbiome-targeted strategies to achieve sustained neuroprotection.

## Acknowledgments.

This work was partly supported by NIH grant R56AG080920 from the National Institute on Aging (NIA), and by the Alzheimer’s Association Research Grant to Promote Diversity. S.S. is supported by the Houston Methodist NeuralCODR Fellowship program, through the generosity of Paula and Rusty Walter and Walter Oil & Gas Corp. A.M. is supported by a training fellowship from the Gulf Coast Consortia on the NLM Training Program in Biomedical Informatics & Data Science (T15LM007093). P.S. was supported by a fellowship from the Coordenação de Aperfeiçoamento de Pessoal de Nível Superior, Brazil (CAPES, Finance Code 001). The authors thank the CPRIT Proteomics and Metabolomics Core Facility (RP210227), NIH (P30 CA125123), and Dan L. Duncan Cancer Center at Baylor College of Medicine and the Pathology and Histology Core at Houston Methodist Research Institute. The authors also thank Cathryn Gunawan and Olivia Chang for their technical help, and Dr. Todd Treangen and Dr. Yunxi Liu from the Department of Computer Science at Rice University for their supervision of the bioinformatic data presented in this study. Figure 1a was done utilizing Biorender.com.

## Data Availability

All raw 16S rRNA sequencing has been deposited into the SRA and is available at BioProject PRJNA1220689.

## Code Availability

All code used within the analysis of the microbiome 16S rRNA sequencing data can be found at https://github.com/villapollab/fmt_ad_no_abx

## Notes

https://github.com/villapollab/fmt_ad_no_abx

### Competing Interest Statement

The authors have declared no competing interest.

